# Cross-talk between the receptor tyrosine kinases AXL and ERBB3 regulates invadopodia formation in melanoma cells

**DOI:** 10.1101/412379

**Authors:** Or-Yam Revach, Oded Sandler, Yardena Samuels, Benjamin Geiger

## Abstract

The invasive phenotype of metastatic cancer cells is accompanied by the formation of actin-rich invadopodia, which adhere to the extracellular matrix, and degrade it. In this study, we explored the role of the tyrosine kinome in the formation of invadopodia in metastatic melanoma cells. Using a microscopy-based siRNA screen, we identified novel invadopodia regulators, the knock-down of which either suppresses (e.g., TYK2, IGFR1, ERBB3, TYRO3, FES, ALK, PTK7) or enhances invadopodia formation and function (e.g., ABL2, AXL, CSK). Particularly intriguing was the discovery that the receptor tyrosine kinase AXL displays a dual regulatory function, manifested by enhancement of invadopodia function upon knock-down or long-term inhibition, as well as following its over-expression. We show here that this apparent contradiction may be attributed to the capacity of AXL to directly stimulate invadopodia; yet its suppression up-regulates the ERBB3 signaling pathway, which consequently activates core invadopodia regulators, and greatly enhances invadopodia function. Bioinformatic analysis of multiple melanoma cells points to an inverse expression pattern of AXL and ERBB3, with the apparent association of high-AXL melanomas, with high expression of invadopodia components and an invasive phenotype. The relevance of these results to melanoma metastasis *in vivo*, and to potential anti-invasion therapy, is discussed.

## Introduction

The morbidity and mortality rates caused by melanoma tumors are primarily associated with the high capacity of these tumors to metastasize, resulting in a 10-year survival rate of 12.5-26% (1). The initial steps of the metastatic process include local invasion of the melanoma cells into the surrounding connective tissue, driven by signaling pathways that stimulate cell migration across tissue barriers; turnover of cell-matrix and cell-cell adhesions; and penetration of the cancerous cells into the vascular and lymphatic systems (2). The migratory and invasive processes are commonly driven by reorganization of the actin cytoskeleton, and the formation of protrusive cell structures such as lamellipodia, filopodia, ruffles, and invadopodia (3).

Invadopodia are actin-rich protrusions of the plasma membrane, which attach to the extracellular matrix (ECM) via integrin receptors, and degrade it (4). These structures are commonly found in metastatic cancer cells, and are believed to promote the dissemination of metastases *in vivo* (5). The local invasion-promoting activity of invadopodia is achieved by coordination of local adhesions, enzymatic degradation of the ECM, and the development of physical protrusive force, generated by actin polymerization, that pushes the plasma membrane outward (4,6). Their proteolytic activity, exerted by secreted and membrane-bound metalloproteinases (MMPs), and serine proteinases (7) distinguishes invadopodia from other migration-promoting membrane protrusions (e.g., lamellipodia, filopodia) or ECM adhesions (e.g., focal adhesions, podosomes) (5).

The temporal sequence of events that drives invadopodia formation and function is not fully understood; yet it has been proposed that this process is initiated by ligand-induced activation of receptor tyrosine kinases (RTKs) (8) that activate Src family kinases (9), and the PI3K-AKT pathway (10). Consequently, TKS5 (5,11), and cortactin (12–14) are phosphorylated, and participate in the early stages of actin machinery assembly in invadopodia (5,6). A few RTKs, such as members of the EGFR family (13,15,16), the DDR family kinases (17), and Tie2 (18), were shown to affect invadopodia formation and function, suggesting that tyrosine phosphorylation is a major regulator of invadopodia formation and function in different cell lines. That said, an understanding of the specific roles of individual kinases in the regulation of invadopodia formation and action remains limited.

We addressed the issue in this study by conducting a microscopy-based siRNA screen, targeting each of the 85 human tyrosine kinases in A375 metastatic melanoma cells, and tested the effect of their knock-down (KD) on invadopodia formation and on ECM degradation. The screen revealed 7 kinases (TYK2, IGFR1, ERBB3, TYRO3, FES, ALK, PTK7), whose suppression decreased invadopodia function, and 3 kinases (ABL2, AXL, CSK) whose suppression increased invadopodia function. Testing the effect of those “hits” on another metastatic melanoma cell line, 63T, confirmed some, but not all, of the hits identified in the original screen, which may be attributed to a different driver mutation in the cells (a BRAF/NRAS mutation in A375/63T cells, respectively) and cell-specific variations in the tyrosine kinome.

In-depth characterization of the role of the receptor tyrosine kinase AXL in invadopodia formation indicated that this kinase has a dual role as an invadopodia regulator, with an intrinsic capacity to stimulate invadopodia; yet its suppression up-regulates an alternative signaling pathway for invadopodia formation: the ERBB3 signaling pathway, which can activate core invadopodia regulators, and greatly enhance invadopodia formation and function. The mechanism underlying the apparent cross-talk between AXL and ERBB3, and its potential relevance to metastatic melanoma therapy, is discussed.

## Results

### Primary screen for novel invadopodia regulators in melanoma cells

The search for invadopodia regulators was conducted by siRNA screening, whereby each of the human tyrosine kinases was knocked down (KD) from the metastatic human melanoma cell line A375 (See Supplementary Material 1). These cells form prominent invadopodia in 30-40% of the cells (19), displaying a moderate capacity to degrade the underlying gelatin matrix, and enabling the quantification of both suppression and enhancement of matrix degradation. Immunolabeling of these cells for phosphotyrosine, together with actin as a marker for invadopodia, revealed extensive localization of tyrosine-phosphorylated residues both within invadopodia cores, and throughout the adhesion rings surrounding them (Fig. 1A; invadopodia are denoted by white arrows). A comparable level of labeling was also detected in focal adhesions located mainly at the cell periphery (Fig. 1A).

**Figure 1:**
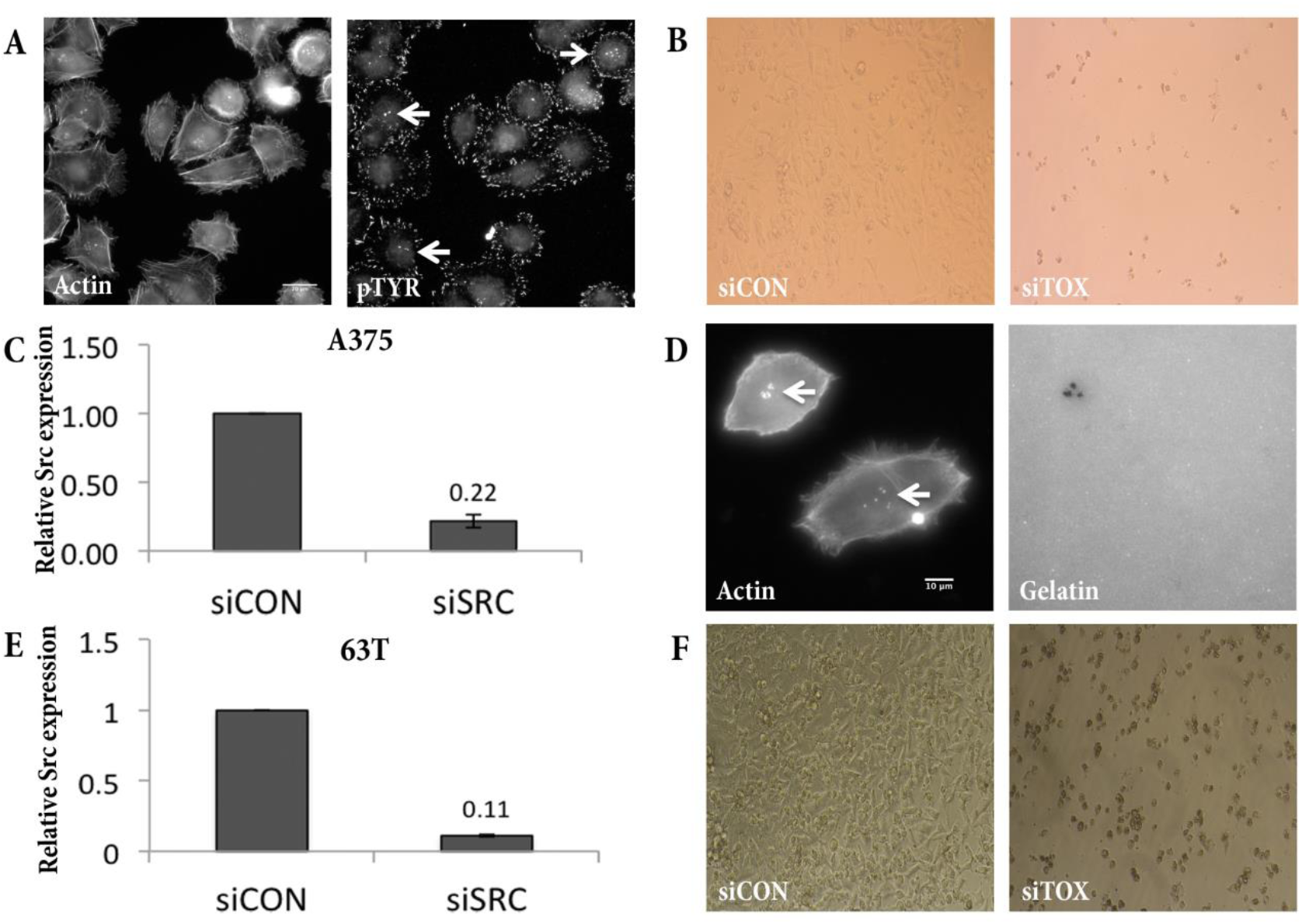
Microscopy-based siRNA screen for tyrosine kinases. (A) A375 cells were cultured on gelatin gel for 3 h and stained for phosphorylated tyrosine and actin. White arrows denote invadopodia structures. Scale bar: 20 μm. (B, F) Representative images from the siCON versus siTOX samples of the screen, used to evaluate transfection efficiency in A375 and 63T cells, respectively. (C, E) Real-time analysis of Src gene expression following transfection under screening conditions, in A375 and 63T cells, respectively; n=3. (D) 63T cells cultured for 3 h on fluorescently labeled gelatin and stained for actin, show that they degrade the matrix (right image), and form actin dots (left image; invadopodia structures are denoted by white arrows). Scale bar: 10 μm.

The siRNA ‘SMARTpool’ library was transfected into A375 cells in a 96-well format, and transfection efficiency was found to be 80-90%, based on the ability of siTOX to kill the cells (Fig. 1B), or quantitative real-time PCR of Src in Src KD cells (Fig. 1C). To assess the effect of the tyrosine kinase KD on invadopodia function, the treated cells were replated on fluorescently-tagged gelatin in 96-well glass-bottomed plates for 6 h, then fixed and stained for actin and TKS5, as invadopodia markers, and DAPI, for cell quantification and assessment of cell toxicity (Supplementary Fig. 1A). Plates were then imaged automatically (see Materials and Methods) and gelatin degradation area per cell (µm^2^) was quantified (Supplementary Fig. 1A and Supplementary Material 1). In order to compare the results of the different experiments, gelatin degradation values were normalized to the control, that was set as 1.

Invadopodia formation was evaluated qualitatively, based on the staining for actin and TKS5 (Table1, Phenotype). For the final hits, quantitative analysis of the percentage of invadopodia-forming cells was performed (Supplementary Fig. 1B, 1C). Only genes whose knock-down resulted in a statistically significant difference (*p* value < 0.05) and led to a >3-fold reduction or a >1.5-fold elevation in gelatin degradation, compared to control, were selected for further characterization (Supplementary Material 1, Screen 2 tab) Altogether, the primary screen was performed three times for the final hits (See full results in Supplementary Material 1).

Of the 85 human tyrosine kinases tested, KD of 7 genes reproducibly reduced gelatin degradation, while 3 enhanced it (Table 1; Fig. 2A-C and Fig. 2B, 2C). Notably, all the “hits” that affected gelatin degradation also induced comparable changes in invadopodia formation (Table 1, Phenotype; and Supplementary Fig. 1B, 1C). KD of each of the three genes causing an elevation in invadopodia function (CSK, AXL and ABL2) resulted in an increase in the percentage of cells forming invadopodia with a high content of TKS5 in their cores (Table 1, Phenotype; Supplementary Fig. 1B, 1C). In contrast, KD of most genes that caused reduction in gelatin degradation resulted in a decrease in invadopodia-forming cells (Table 1, Phenotype; Supplementary Fig. 1B, 1C).

**Table 1:** Invadopodia regulators in A375 cells. Normalized degradation/cell is presented for each gene knock-down in A375 cells. *P*-value for each gene is presented. Three repeats; n> 100 for each sample. The phenotype caused by knockdown of each of the genes, visualized by actin and TKS5 staining and evaluated in a qualitative fashion, is described under the Phenotype heading.

| Gene knockdown | Degradation/cell (normalized) | *p*-value | Phenotype |
| --- | --- | --- | --- |
| TYK2 | 0.19 | 4.03E-04 | less invadopodia |
| IGF1R | 0.38 | 9.24E-05 |  |
| TYRO3 | 0.21 | 1.54E-03 | less invadopodia |
| FES | 0.17 | 4.59E-03 | less invadopodia |
| ALK | 0.26 | 3.88E-11 | more structures, non mature |
| ERBB3 | 0.32 | 9.79E-05 | small stuctures, no TKS5 |
| PTK7 | 0.34 | 1.92E-03 | less invadopodia |
| control | 1.00 |  |  |
| ABL2 | 2.27 | 2.31E-10 | more invadopodia |
| AXL | 2.84 | 5.60E-03 | more invadopodia |
| CSK | 8.31 | 6.77E-03 | more invadopodia |

**Figure 2:**
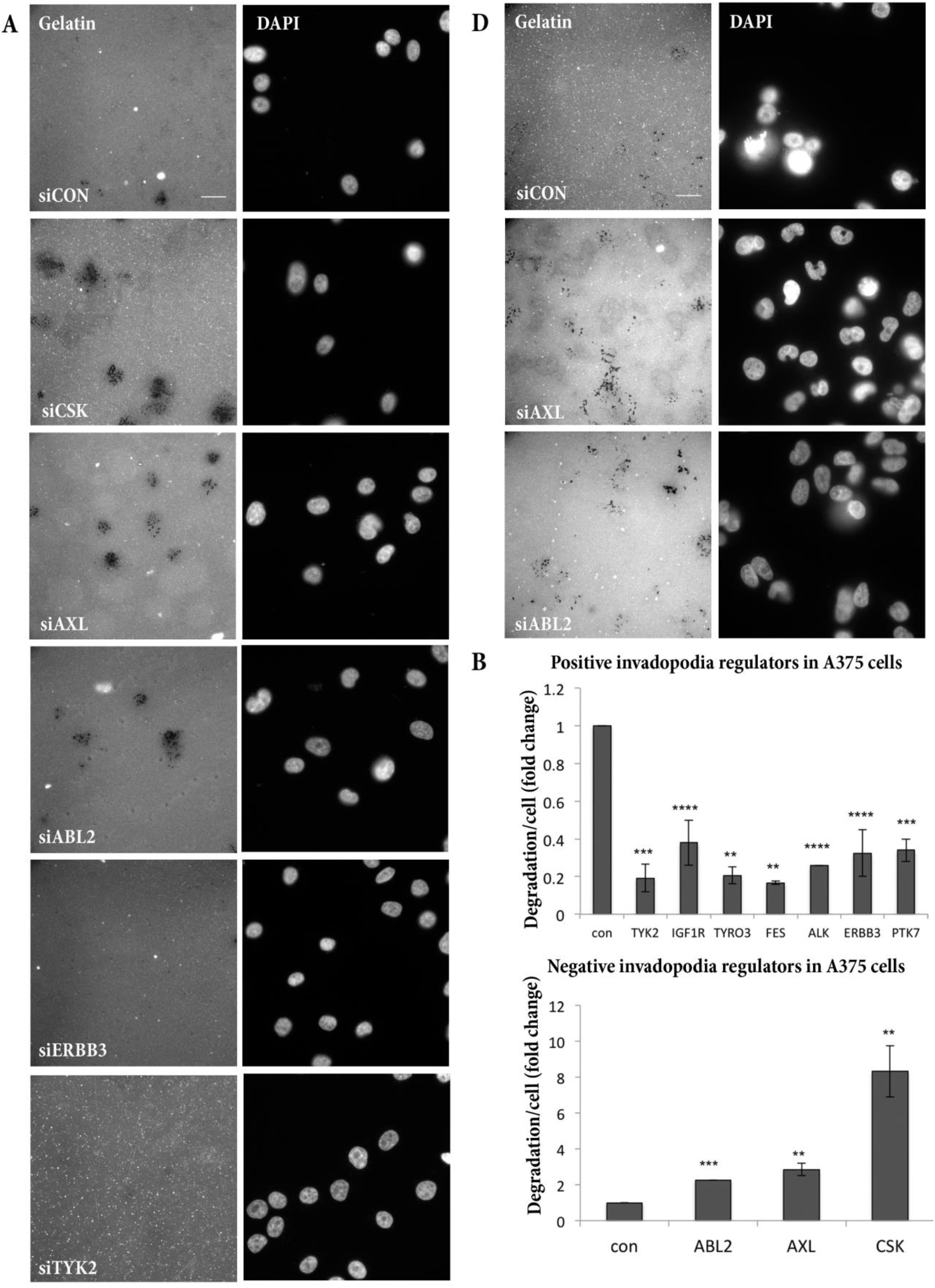
Screen results for invadopodia regulators of the tyrosine kinase family. (A) Representative images of A375 cells from the screen. Cells were cultured for 6 h on fluorescently labeled gelatin, and stained with DAPI. Scale bar: 20 μm. (B, C) Quantification of normalized degradation/cell for the final hits of the screen (invadopodia assay) in A375 cells. C shows genes that caused a reduction in invadopodia function upon KD, and D shows genes that caused an elevation of invadopodia function upon KD. Three repeats; n>100 in each sample. (D) Same as in A, for 63T cells. Scale bar: 20 μm.

It is noteworthy that some of the invadopodia-regulating genes identified in this screen were previously shown to affect invadopodia formation or function, including CSK (20), which inhibits the activity of Src family kinases (21), and upon knock-down, causes a dramatic elevation in invadopodia function. PTK7 and ERBB3 were shown to be positive invadopodia regulators in other cellular systems (22,23), and ABL2 was found in our screen of melanoma cell lines to be a negative regulator, while in breast cancer cells, it was shown to positively regulate invadopodia (13,24).

Given the potential functional redundancy between tyrosine kinases, which may vary in different cells, we further tested the effectiveness of all the screen’s “hits” on another metastatic, patient-derived melanoma cell line, 63T, which differs from A375 cells in its genetic background (a BRAF/NRAS mutant in A375/63T cells, respectively) (25). The 63T cells display conspicuous invadopodia in 20-30% of the cells with extensive matrix degradation capacity (Fig. 1D), and are suitable for siRNA transfection (Fig. 1E, 1F). Testing the effect of siRNA KD of the A375 screen hits on 63T cells (see Supplementary Material 1), indicated that only the KD of FES, one of the 7 suppressive hits obtained with A375 cells, also induced a reduction in invadopodia function in 63T cells. Conversely, the 3 whose KD enhanced invadopodia in A375 cells (CSK, AXL and ABL2), had a similar effect on 63T cells (Table 2; Fig. 2D; and Supplementary Fig. 1D).

**Table 2:** Invadopodia regulators in 63T cells. Same as Table 1, for 63T cells. Two repeats; n> 100 for each sample.

| Gene knockdown | Degradation/cell (normalized) | *p*-value |
| --- | --- | --- |
| FES | 0.10 | 2.76E-03 |
| siCON | 1.00 |  |
| AXL | 3.95 | 2.16E-04 |
| ABL2 | 5.02 | 1.61E-07 |
| CSK | 11.58 | 1.31E-15 |

AXL is a receptor tyrosine kinase, associated with an invasive phenotype in multiple cancers, including melanoma (26,27). In addition, AXL was shown to be involved in drug resistance mechanisms in melanoma and other cancers (28,29)

Intrigued by the unexpected enhancement of invadopodia following the KD of AXL, and its novel association with invadopodia regulation, we chose to focus our study on the mechanism underlying the effect of this kinase on invadopodia formation and function.

### The AXL receptor tyrosine kinase regulates invadopodia in melanoma cells

To validate the specificity of the augmentation of invadopodia prominence and matrix degradation upon AXL KD, we tested each of the four duplexes present in the siRNA ‘SMARTpool’ in both cell lines, and confirmed that 3 out of 4 siRNA duplexes display an elevated degradation effect similar to that seen in siAXL siRNA pool (Supplementary Fig. 2A, 2B). Further quantification of the effect indicated that upon KD of AXL, normalized gelatin degradation levels (degraded area/cell) increased by a factor of 7 (in A375 cells) or of 4 (63T cells) (Fig. 3A-C), and the percentage of invadopodia-forming cells in both cell lines was comparably elevated (Fig. 3E, 3F, respectively). AXL suppression efficiency was up to 90% determined by Western blot (Fig. 3D) and real-time PCR (Supplementary Fig. 2C, 2D). A similar effect of elevated invadopodia function was obtained when cells were stably infected with shAXL (Supplementary Fig. 2E, 2F), and upon siRNA transfection in two other melanoma cell lines, WM793 and A2508, that form invadopodia (Supplementary Fig. 2G-I).

**Figure 3:**
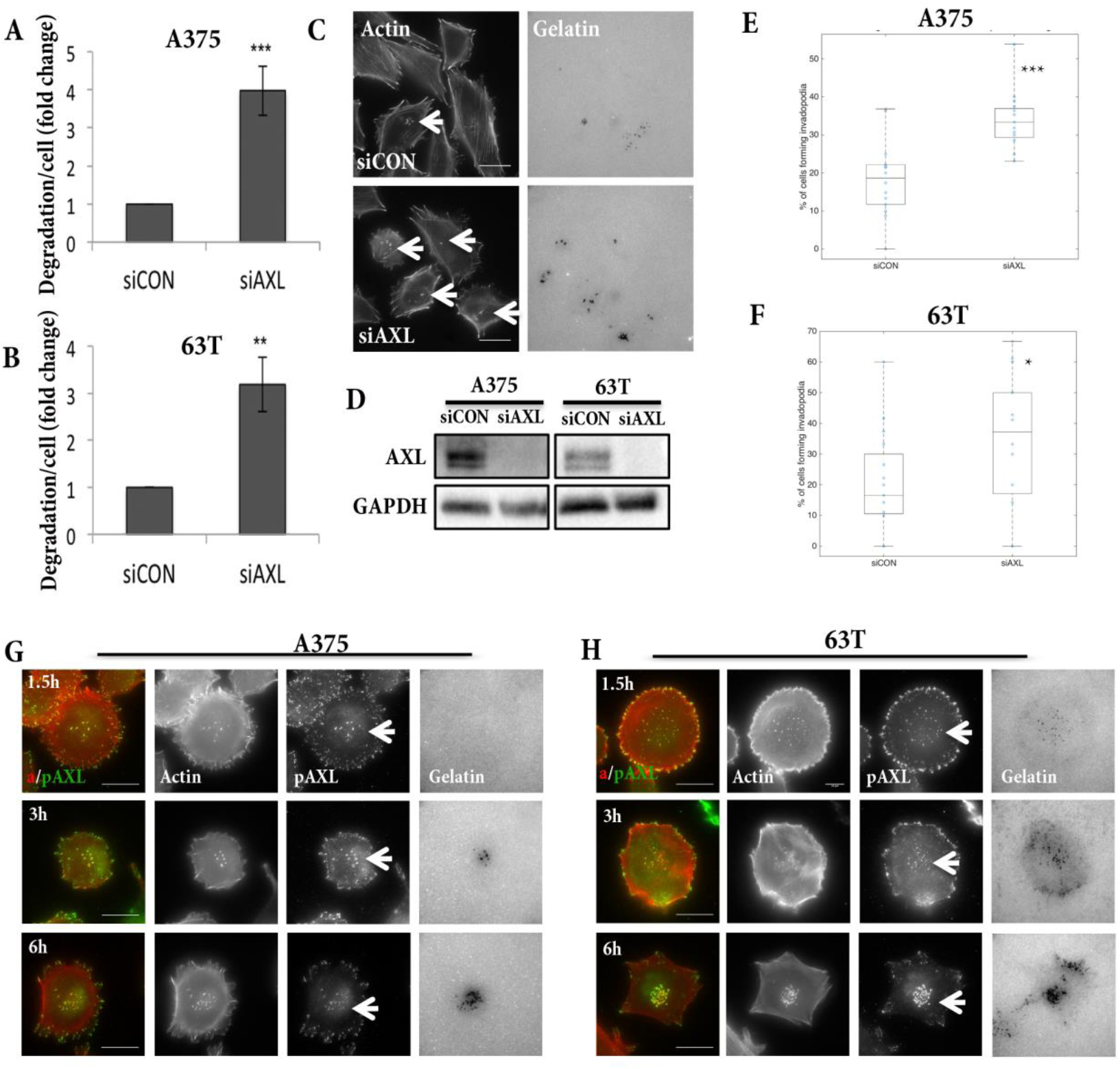
AXL is localized to invadopodia, and regulates their function. **(A, B)** Invadopodia assay of siCON versus siAXL in A375 and 63T cells, respectively. Cells were cultured for 6 h. Ten repeats for each cell; n>100 in each sample. (C) Representative image of A375 siCON and siAXL cells. Scale bar: 20 μm. (D) Western blot analysis of AXL protein in A375 and 63T cells upon AXL knock-down. GAPDH was used as a loading control. (E, F) Quantification of invadopodia-forming cells (expressed as a percentage) in A375 and 63T cells, respectively. Cells were cultured on gelatin gel for 6 h. Three repeats; n>100 in each sample. (G, H) A375 (G) or 63T cells (H) were cultured on fluorescently labeled gelatin for 1.5 h, 3 h, and 6 h, and stained for actin and phosphorylated AXL. AXL localized to invadopodia cores from the early stages of their formation (1.5 h), up to the mature stage (6 h). AXL also localized to focal adhesions. Invadopodia structures are denoted by white arrows. Scale bar: 20 μm.

The role of invadopodia in elevated matrix degradation induced by AXL suppression was confirmed by double knockdown of both AXL and TKS5 (an invadopodia-specific marker), which eliminated the elevation in matrix degradation (Supplementary Fig. 2J, 2K). Finally, localization of phosphorylated AXL in A375 and 63T melanoma cells, as seen by immunofluorescence microscopy, pointed to a strong association of AXL with invadopodia cores at various time points following cell culturing and invadopodia formation (Fig. 3G, 3H). Taken together, these results suggest that AXL plays an important role in the regulation of invadopodia in melanoma cells.

### AXL is a dual-function regulator of invadopodia, with both suppressive and stimulatory activity

Further characterization of the molecular mechanisms underlying the role of AXL in invadopodia formation and function indicated that AXL might have a dual regulatory effect, both negative and positive. First, we tested the effect of AXL overexpression on invadopodia function (e.g., gelatin degradation) in A375 cells (Fig. 4A, 4B) or in 63T cells (Fig. 4C). The over-expression of wild-type AXL, but not of its kinase-dead derivative, or empty vector, caused a significant elevation in invadopodia function, suggesting that kinase activity is required for the positive regulation of invadopodia (Fig. 4A-C). The transfection resulted in a marked elevation in total AXL levels (Supplementary Fig. 3A). The elevation in gelatin degradation also correlated with the ability of these cells to invade through a transwell filter *in vitro* (Supplementary Fig. 3B, 3C).

**Figure 4:**
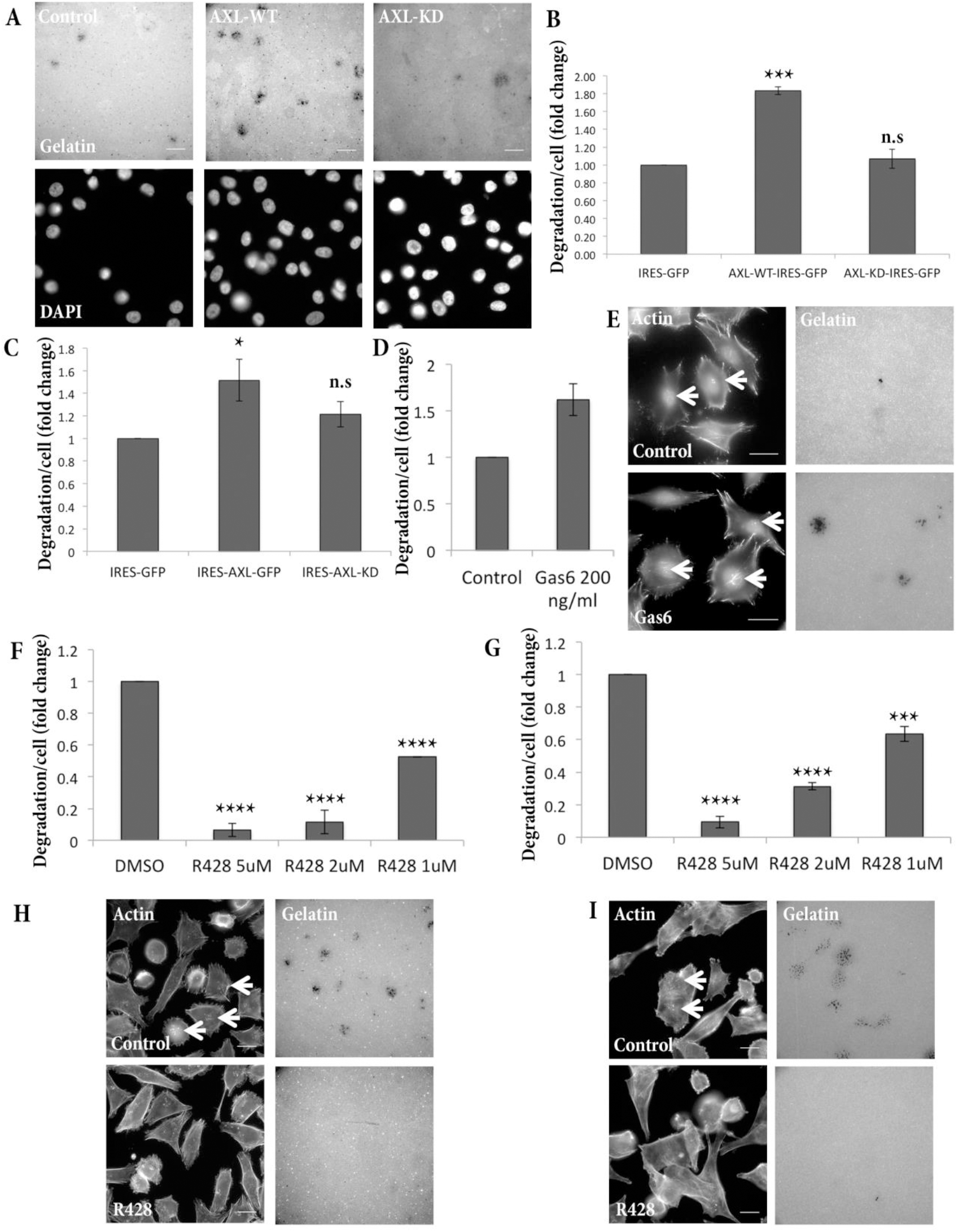
AXL is a positive regulator of invadopodia in melanoma cells. Invadopodia assay of A375 cells (A, B) or 63T cells (C) overexpressing AXL-WT, AXL kinase-dead (AXL-KD), or control plasmid, cultured on fluorescently labeled gelatin for 6 h, and stained for DAPI. Representative images in A; quantification in B and C. Three repeats for each cell type; n>100 in each sample. Scale bar: 20 μm. (D, E) Invadopodia assay of control or 200 ng/ml Gas6-treated cells cultured on fluorescently labeled gelatin for 6 h. Gas6 effective concentrations were varied between experiments, and ranged between 200-1000 ng/ml. Four repeats, n>100 in each sample. Scale bar: 20 μm. (F, G) Invadopodia assay of A375 (F) and 63T cells (G) cultured on fluorescently labeled gelatin for 6 h with DMSO or the AXL inhibitor R428 (in concentrations of 5μM, 2μM, or 1μM). Three repeats for each cell type; n>100 in each sample. (H, I) Representative images of A375 cells in 2μM concentrations (H) or 63T cells in 5μM concentrations (I), stained for actin and cultured on fluorescently labeled gelatin. Invadopodia structures are denoted by white arrows. Scale bar: 20 μm.

Additional support for the positive role of AXL in invadopodia formation and function in A375 or 63T melanoma cells was obtained through elevated levels of invadopodia, following treatment with the AXL ligand Gas6 (Fig. 4D and Supplementary Fig. 3D), and the suppression of invadopodia in both cell types by treatment with the AXL inhibitor R428 [Fig. 4F, 4H and 4G, 4I]. Invadopodia inhibition by the AXL inhibitor was also observed in two additional, patient-derived melanoma cells lines, 76T and 104T, as well as, in 4T1 murine breast cancer cells that form invadopodia (Supplementary Fig. 3E, 3F). Elevated levels of invadopodia function upon AXL knock-down was specific to melanoma cells: when AXL was knocked down in MDA-231 breast cancer cells, invadopodia function was decreased by factor of two (Supplementary Fig. 3G, 3H), corroborating our observation that AXL acts as a general positive invadopodia regulator.

### ERRB3 activation is up-regulated upon AXL suppression, leading to enhanced invadopodia function

Given existing evidence that AXL promotes invasion and migration in a variety of cancer cells (30–32), together with our current findings, we hypothesized that an alternative invasion-promoting pathway may be activated following AXL inhibition by siRNA, leading to an elevation in invadopodia formation. Furthermore, this alternative pathway is not activated in instances of short-term inhibition of AXL (Fig. 4F-I).

Cross-talk between AXL and EGFR family members was previously proposed by studies in several cancer systems as a potential mechanism for drug resistance, usually in the form of AXL activation following EGFR inhibition (33–35). Moreover, in breast cancer cells, the opposite process was found, when prolonged inhibition of AXL leads to an increase in ERBB3 expression and phosphorylation (36). This phenotype was not linked to invasion or invadopodia. Moreover, to our knowledge, no such interplay between AXL and ERBB3 was documented in melanoma cells.

When we analyzed, in A375 cells, the levels of ERBB3 activation under conditions of AXL KD, we found an increase in total ERBB3 levels, as well as in ERBB3 phosphorylation (Fig. 5A). RNA levels of ERBB3 were only slightly elevated following AXL inhibition (Supplementary Fig. 3I), suggesting that the up-regulation of ERBB3 is primarily post-transcriptional. Moreover, Src, Cortactin and AKT-activating phosphorylation levels were also elevated under those same conditions (Fig. 5B, 5C, and 5D, respectively). Activation of these components is an indication of more active invadopodia, and higher levels of actin polymerization (10,12,14,37). Indeed, the SRC and PI3K-AKT pathways can potentially be activated directly by ERBB3 (38,39) and cortactin is a downstream target of Src in invadopodia formation (5,12,13). These results suggest that AXL inhibition activates the ERBB3 pathway in melanoma cells, and this activation may lead to an increase in invadopodia formation.

**Figure 5:**
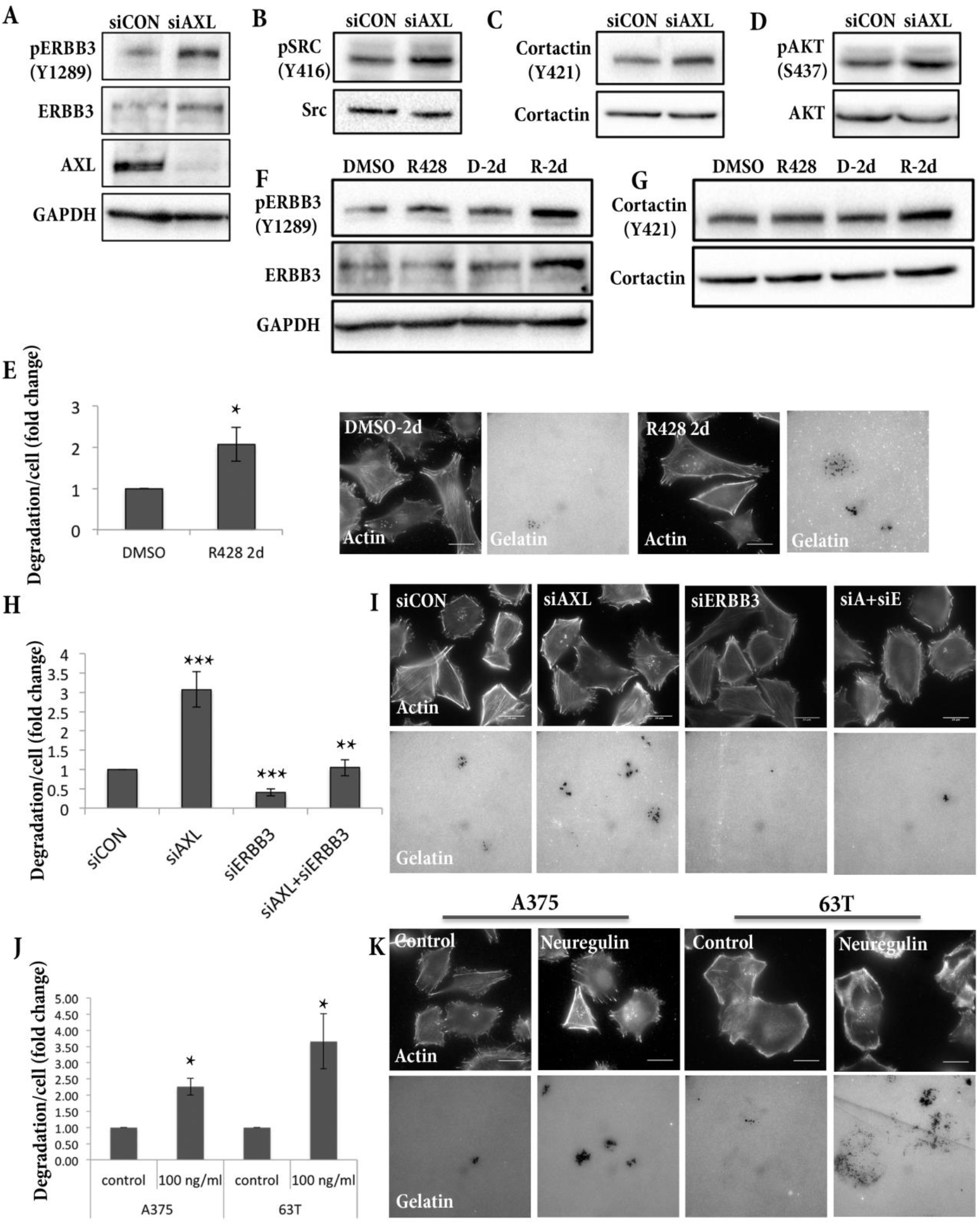
The ERBB3 pathway is activated when AXL is inhibited for prolonged periods. (A) Western blot of A375 cells, showing ERBB3 phosphorylation (Y1289) and total ERBB3 protein in siAXL versus siCON cells. AXL levels are presented, as well as that of the GAPDH loading control. (B) Western blot of A375 cells, showing Src phosphorylation (Y416) and total Src protein levels in siAXL versus siCON cells. (C) Western blot of A375 cells, showing cortactin phosphorylation (Y421) and total cortactin protein levels in siAXL versus siCON cells. (D) Western blot of A375 cells, showing AKT (S437) phosphorylation and total AKT protein levels in siAXL versus siCON cells. For A-D, n=3. (E) Invadopodia assay of A375 cells incubated for 2 days with 1μM R428, AXL inhibitor, or DMSO. After 48 h, cells were replated on fluorescently labeled gelatin for 6 h for the invadopodia assay. Invadopodia are denoted by white arrows. Four repeats; n>100 in each sample. (F) Western blot of A375 cells, showing ERBB3 phosphorylation (Y1289) and total protein in DMSO (D) versus R428 (1μM) short-term treatment (6 h) and DMSO (D-2d) versus R428-2d (1μM) prolonged treatment (2 days). GAPDH as loading control (G) Western blot of A375 cells, showing cortactin phosphorylation (Y421) and total protein in DMSO (D) versus R428 (1μM) short-term treatment (6 h) and DMSO (D-2d) versus R428-2d (1μM) prolonged treatment (2 days). For F and G, n=3. (H, I) Invadopodia assay of A375 cells with KD of AXL, ERBB3, AXL+ERBB3, and control cells cultured on fluorescently labeled gelatin for 6 h. Six repeats; n>100 in each sample. (J, K) Invadopodia assay of A375 or 63T cells (two repeats or three repeats, respectively; n>100 in each sample) cultured on fluorescently labeled gelatin for 6 h with 100 ng/ml neuregulin, or vehicle.

Consistently, invadopodia function was also elevated following long-term (2 days’) inhibition of AXL by the R428 inhibitor, comparable to that observed following AXL siRNA-mediated KD (Fig. 5E), and in contrast with short-term (6h) exposure to the inhibitor, which readily blocks invadopodia formation (Fig. 4F-4I). Moreover, prolonged AXL inhibition using R428 results in higher levels of total and phosphorylated ERBB3 (Fig. 5F, 2 right lanes). Notably, no such elevation was observed following short-term (6h) treatment with the inhibitor (Fig. 5F, 2 left lanes). Cortactin phosphorylation was also elevated under prolonged AXL inhibition using R428 (Fig. 5G, 2 right lanes), and not under short-term treatment (Fig. 5G, 2 left lanes).

The notion that the elevation in invadopodia function in siAXL KD cells depends on ERBB3 activation was further tested by double-KD experiments, which confirmed that invadopodia elevation by AXL knock-down does not take place in the absence of ERBB3 (Fig. 5H and 5I). In accordance with the siRNA screen, KD of ERBB3 alone led to partial suppression of invadopodia function (Fig. 5H; Fig. 2B; Table 1) Finally, Neuregulin, the ERBB3 ligand, stimulated elevated invadopodia function in both cell types (Fig. 5J, 5K).

Our findings suggest that when AXL is suppressed or inhibited for prolonged periods, the ERBB3 pathway, an alternative, highly potent signaling pathway for invadopodia formation, is activated, leading to increased invadopodia formation and function.

### AXL expression exhibits an expression pattern inverse to that of ERBB3 and MITF in melanoma cells

To test whether the interplay between AXL and ERBB3 constitutes a common feature of melanoma cells, we analyzed the expression levels of AXL and ERBB3 in 62 melanoma cell lines from the *CCLE* database (https://portals.broadinstitute.org/ccle). MITF and three of its downstream targets were added to the analysis, due to the known inverse correlation of AXL and MITF expression levels in melanoma, and the relevance of MITF/AXL ratio to melanoma state, progression and drug resistance (40,41).

On the CCLE database-derived expression Z-score values, we applied hierarchical clustering, and four groups of cell lines were manually defined (Figure 6A). Two of the clusters (Cluster 1 and Cluster 2; a total of 31 cell lines) encompassing 50% of the dataset, showed a distinct, inverse expression pattern between AXL and ERBB3 (p-value <0.0006; see Supplementary Material 3). Moreover, ERBB3 expression levels were positively correlated with the levels of MITF and its downstream targets (Fig. 6A, Clusters 1 and 2). Clusters 3 and 4 showed different expression patterns: either all three genes (AXL, ERBB3 and MITF) were high (Fig. 6A, cluster 3); or both ERBB3 and AXL were high, but that of MITF was low (Fig. 6A, Cluster 4). This analysis demonstrates that in many cases of melanoma (50%, in our dataset), the expression patterns of AXL and ERBB3 are mutually exclusive, suggesting that they act on alternative oncogenic pathways, as we see in the context of invadopodia regulation (Fig. 5).

**Figure 6:**
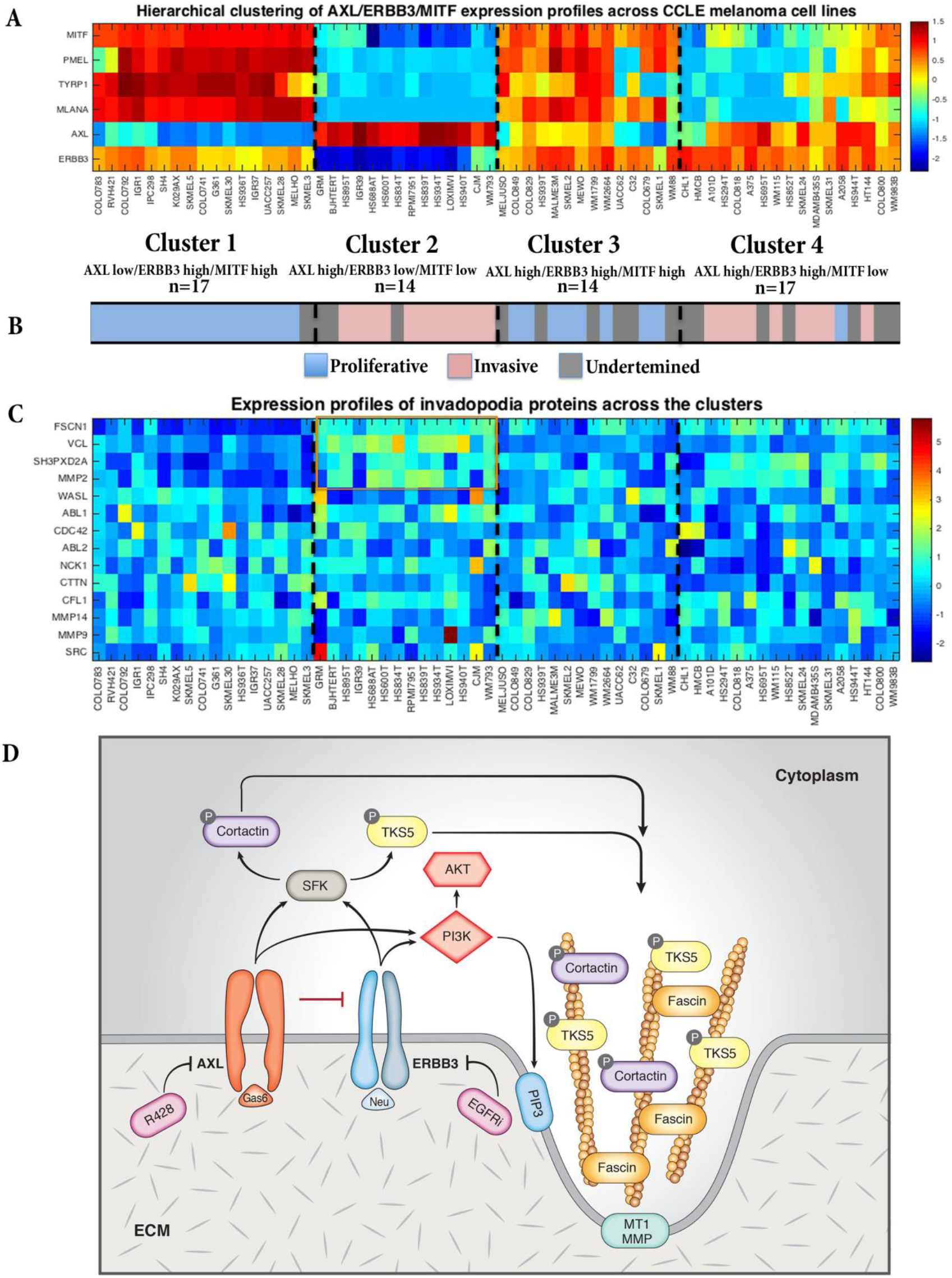
AXL negatively correlates with ERBB3 in 50% of melanomas. (A) Hierarchical clustering by Euclidian distance was applied on the Z-score values of AXL, ERBB3, MITF, and three MITF targets: PMEL, TYRP1, and MELANA, from 62 melanoma cell lines of the *CCLE* Cell Line dataset. Four clusters were selected manually. AXL and ERBB3 are significantly negatively correlated (for *p*-values, see Supplementary Material 3) (B) Prediction of cell phenotype based on the MPSE (Melanoma Phenotype-Specific Expression) signature (42,43). (C) Analysis of Z-score values of 14 core invadopodia proteins among the four clusters described in A. *P*-values for the differences of the proteins in Clusters 1 and 2 are presented in Supplementary Material 3. (D) Proposed model for AXL-ERBB3 interplay in invadopodia regulation. AXL and ERBB3 can both positively regulate invadopodia, depending on their initial balance in the cell. Each of them is activated by its ligand (AXL by Gas6; ERBB3 by neuregulin). A one-directional interplay exists between the two receptors. Once AXL is subjected to short-term inhibition (6 h) by the inhibitor, invadopodia function is inhibited; while when AXL is subjected to prolonged inhibition (2 days) by siRNA or inhibitor treatment, ERBB3 can be activated, its protein levels are elevated, and a corresponding elevation in invadopodia function is seen. Both AXL and ERBB3 can activate PI3K and SFK, leading to phosphorylation of TKS5 and cortactin, that integrate into the invadopodia core, and initiate their formation. Further actin polymerization and actin bundling by Fascin occurs, eventually resulting in the secretion of MMPs, and matrix remodeling.

Next, we classified the cells as proliferative or invasive, based on their melanoma phenotype-specific expression (MPSE) signature (42,43). MPSE predicts whether the cell phenotype will be proliferative or invasive, based on the expression of a set of genes. Cluster 1 (AXL low/ERBB3 high) is enriched with proliferative cell lines (Fig. 6B, lower bar), whereas Cluster 2 (AXL high/ERRB3 low), is dominated by an invasive phenotype (Fig. 6B, lower bar).

Consistently, Cluster 2 exhibits a significantly higher expression of invadopodia components [e.g., TKS5 (SH3PXD2A), Fascin, MMP2, ABL1, and vinculin; see Fig. 6C, and Supplementary Material 3 for *p-*values].

To gain an overview of the contribution of the tyrosine kinome to melanoma invasion and invadopodia formation, we analyzed the expression patterns of all tyrosine kinases, and their distribution among the 4 clusters (Supplementary Fig. 4). A group of 13 tyrosine kinases were co-expressed with high AXL/low ERRB3 levels (Cluster 2; see Supplementary Material 3); among these, PDGFRA and PDGFRB, EGFR and FGFR1. In contrast, another group of 11 tyrosine kinases negatively co-expressed with high ERRB3/low AXL (Cluster 1; see Supplementary Material 3); among them, two other members of the TAM receptor family MER and TYRO3, as well as Syk and CSK (Supplementary Fig. 4). Other tyrosine kinases were not differentially expressed among the clusters. This analysis further supports the notion of two alternative pathways that regulates cell invasion in melanoma.

## Discussion

In this study, we describe the molecular cross-talk between different tyrosine kinases involved in the regulation of invadopodia formation and function in melanoma cells. Using a comprehensive siRNA screen, we identified several novel positive invadopodia regulators in A375 metastatic melanoma cells, including TYK2, IGF1R, TYRO3, FES, ALK, ERBB3 and PTK7 (Fig. 2A, 2B; Table 1). In addition, we found 3 kinases (CSK, ABL2 and AXL) whose KD up-regulated invadopodia formation and function (Fig. 2A, 2C; Table 1). A few of the hits (CSK, ABL2, ERBB3 and PTK7) were found to regulate invadopodia in other cellular systems (5,13,21,23,44). CSK is, indeed, a negative regulator of invadopodia, as it inhibits SRC family kinases (SKF) (5,21). Src kinases are known to play an essential role in invadopodia formation by phosphorylating and activating key structural components of invadopodia, including, TKS5 and cortactin (12).

It is noteworthy that while KD of CSK, which phosphorylates multiple SFKs, elevates invadopodia formation (5), knock-down of individual SFKs (including Src) had a limited effect on invadopodia formation (Supplementary Material 1). This phenotype is attributable to the compensatory activity of other SFKs, which are prominently expressed in melanoma tumors (45,46).

Similarly, functional redundancy between tyrosine kinases that regulate invadopodia was also implied by marked differences between the profiles of invadopodia modulators in A375 and in another melanoma cell line, 63T (Fig. 2D; Supplementary Fig. 1D; Table 2). Specifically, only 4 genes out of 10 caused similar effects on invadopodia in 63T cells. (See Table 1, Table 2 and Supplementary Fig. 4). These differences may also be attributed to the different driver mutations in the cells, as A375 is a BRAF mutant, while 63T is an NRAS mutant. Clarification of these cellular variations is required for a deeper understanding of invadopodia formation, and for their roles in melanoma cell metastasis.

In this comprehensive analysis, we found that the AXL receptor tyrosine kinase acts as a dual function regulator of invadopodia in both the A375 and 63T melanoma cell lines. AXL is known to be associated with an invasive phenotype in multiple cancers, including melanoma (26,27,47). In addition, AXL was shown to be involved in drug resistance mechanisms in melanoma and other cancers (28,29,48). Our novel finding that AXL is involved in the regulation of invadopodia formation and function (Fig. 3; Fig. 4) adds another dimension to our understanding of how AXL contributes to the development of these aggressive phenotypes.

The dual role of AXL is manifested in the increase of invadopodia formation and function upon AXL KD in 4 different melanoma cells lines (A375, 63T, WM793, and A2058) (Fig. 3A-3F; Supplementary Fig. 2G, 2H). On the other hand, we demonstrate the positive role of AXL in invadopodia formation and function, based on a series of experiments involving two melanoma cell lines (A375 and 63T). AXL localizes specifically to invadopodia structures at various stages of their formation (Fig. 3G, 3H) and causes an increase in invadopodia activity following overexpression of WT-AXL, but not its kinase-dead mutant (Fig. 4A-4C). Consequently, the capacity of these cells to invade in a Matrigel trans-well invasion assay was increased as well (Supplementary Fig. 3B, 3C). Activation of AXL by its ligand, Gas6, increased invadopodia function in both cell lines (Fig. 4D, 4E and Supplementary Fig. 3), while short-term (6 hr) treatment of cells with the AXL inhibitor R428 was found to block invadopodia function in 4 melanoma cell lines (Figure 4F–4I; Supplementary Fig. 3E, 3F), as well as in 4T1 breast cancer cells (Supplementary Fig. 3E, 3F). Finally, knock-down of AXL in the metastatic breast cancer cell line MDA-231 caused a significant reduction in invadopodia function (Supplementary Fig. 3G, 3H).

Further experiments clarified the dual role of AXL in the regulation of invadopodia. AXL can potently promote invadopodia formation (Fig. 4). Upon prolonged AXL inhibition (knock-down or treatment with AXL inhibitor for 2 days), ERBB3 expression is increased and becomes hyper-phosphorylated (Fig. 5A, 5F), leading to an increase in invadopodia function (Fig. 3A, 3B, 5E). ERBB3 activation and increased invadopodia function was accompanied by increased phosphorylation of key components and regulators of invadopodia, including SRC, AKT and cortactin (Fig. 5B-5D, 5G). The role for ERBB3 in invadopodia function is also supported by the fact that ERBB3 was one of the hits in the A375 screen described above (Fig. 2A, 2B; Table 1), and by the augmentation of invadopodia formation upon treatment of A375 or 63T cells with the ERBB3 ligand neuregulin (Fig. 5J, 5K). The fact that only prolonged inhibition of AXL, and not short-term inhibition (6h) leads to elevated ERBB3 levels (Fig. 5F) and consequently, to increased invadopodia formation (Fig. 5E), suggests that the compensatory mechanism is regulated either at the transcriptional level, or at a slow, post-translational process that affects ERBB3 stability or turnover. Notably, similar reciprocal cross-talk between AXL and ERBB3 was reported in breast cancer cells, where prolonged (1-2 days’) inhibition of AXL by inhibitor or siRNA treatment led to an elevation in ERBB3 phosphorylation and activity (36).

Is the ability of ERBB3 to compensate for AXL loss in invadopodia regulation unique? We can’t exclude the possibility that other TKs can compensate for the loss of AXL in invadopodia formation; yet double knock-down of both AXL and ERBB3 is sufficient to abrogate the increase in invadopodia formation. (Fig. 5H, 5I). Notably, the abrogation of invadopodia formation was more conspicuous following siERBB3 KD only, in comparison to double KD of both siERBB3 and siAXL (*p* value=0.01). This observation suggests that AXL inhibits ERBB3 depended invadopodia activation and, upon long-term inhibition of AXL, this inhibition is unleashed, resulting in increased invadopodia function (Fig. 3A, 3B and Fig. 5E).

Examination of AXL/ ERBB3 expression levels in multiple melanoma cells, in the *CCLE* database shows a significant inverse expression patterns of in about 50% of the cells (Fig. 6A, Clusters 1 and 2). Interestingly, ERBB3 levels positively correlate with MITF expression levels, and with its three main downstream targets (Fig. 6A, Clusters 1 and 2). MITF has been shown to negatively correlate with AXL in melanoma cells (29,40). MITF levels are also known to affect melanoma progression, and fine-tuning of its expression leads to various physiological and pathological manifestations, including cell differentiation, proliferation, apoptosis and invasion (40,41). Furthermore, the balance between MITF and AXL can change cell fate, invasiveness and drug resistance (40,41). For example, AXL-high / MITF-low cells tend toward a migratory and invasive phenotype (49).

The AXL/ ERBB3 mutually exclusive expression patterns suggests two modes of action for invadopodia regulation in melanoma (Fig. 6D, Proposed Model). One is governd by AXL which is activated by Gas6 (50), and can promote cell invasion and migration (31,32,47), cytoskeletal remodeling (51–53) and invadopodia formation and function (our results, Fig. 4). This can be achieved by activation, of invadopodia-associated proteins (e.g., TKS5 and cortactin) by Src family kinases and the PI3K pathway (13,14,37,54) see also Fig. 6D, Proposed Model. Indeed, high expression levels of core invadopodia components (4) such as TKS5, MMP2, ABL1, Fascin and vinculin are correlating with high AXL levels (Fig. 6C, Cluster 2); as well as an invasive phenotype (Fig. 6A, 6B, Cluster 2).

The second invadopodia-stimulating pathway is induced by ERRB3, which is predominantly correlateing with a proliferative phenotype (Fig. 6B), yet, upon suppression of AXL it takes the lead in promoting invadopodia formation and function, most likely, by activating similar downstream signals (Fig. 5; Fig. 6D, Proposed Model).

The process of cell invasion includes differential regulation of the migratory and invasive capacities of the cells (2). A finely-tuned balance between the invadopodia regulators AXL and ERBB3 is necessary for efficient cancer dissemination, as well as in acquisition of drug resistance in melanoma (41,55). Our new insights into the unique abilities of AXL and ERBB3 to regulate invadopodia function and cell invasion, together with a deeper understanding of the kinome network, and deciphering of the cross-talk between other invadopodia regulators, will facilitate the design of an effective combinatorial therapy, based on the simultaneous supporession of both kinases, that may prevent invadopodia-mediated melanoma cell invasion, and potentially reduce drug resistance.

## Acknowledgments

We thank Prof. Yosef Yarden (Weizmann Institute of Science, Israel) for providing reagents for ERRB3 detection, and scientific advice regarding ERBB3. We thank Dr. Aaron Meyer (Massachusetts Institute of Technology, USA) for his generous gift of AXL expression plasmids. We thank Dr. Itay Tirosh (Weizmann Institute of Science, Israel), Prof. Daniel Peeper and Dr. Oscar Krijgsman (Netherlands Cancer Institute, Amsterdam, Netherlands) and Prof. Tamar Geiger (Tel-Aviv University, Israel) for insightful discussions and scientific advice regarding AXL/ERBB3 cross-talk in melanoma.

Grants: Israel Ministry of Science (IMOS) French program Grant no.3-13024, and the Israel Science Foundation Grant no. 2749/17.

Benjamin Geiger is the incumbent of the Erwin Neter Professorial Chair in Cell and Tumor Biology.

## Author contributions

O-Y. R. performed all the experiments, and contributed to their design, and to the writing of the manuscript. O.S. assisted in analyzing the gene expression patterns. Y.S provided the melanoma cell lines and relevant reagents (See Materials and Methods), as well as scientific contributions regarding melanoma biology and gene expression analysis. B.G. contributed to the design of the experiments, examination of the results, and to the writing of the manuscript.

## Materials and Methods

### Antibodies and reagents

R428 inhibitor and Gas6 were purchased from R&D Systems (Minneapolis, MN, USA). R428 was used at concentrations of 1-5 μM, depending on the cell type. Gas6 was used at a range of concentrations, 200ng-1,000 ng/ml, depending on the cell type. Effective concentrations of Gas6 varied between experiments. Neuregulin was purchased from PeproTech (Rocky Hill, NJ, USA) and was used in concentrations of 100 ng/ml. Puromycin was used for selection of infected cells at concentrations of 3μg/ml (Sigma St. Louis, MO, USA).

Antibodies used in this study included: Mouse monoclonal anti-phosphorylated tyrosine is a self-prepared soup Rabbit polyclonal anti p-AXL(Y779) was purchased from R&D Systems. Rabbit monoclonal antibody anti p-ERBB3 (Y1289) # 4791; rabbit polyclonal antibody anti-Cortactin # 3503; and rabbit polyclonal antibody p-SRC (Y416) # 2101 were ordered from Cell Signaling (Danvers, MA, USA). Mouse monoclonal anti-ERBB3, SC-7390; rabbit polyclonal anti-TKS5; goat polyclonal anti-AXL, SC-1096; rabbit polyclonal anti-SRC, SC-19; rabbit polyclonal anti-AKT, SC-8312 and rabbit polyclonal anti-p-AKT(s437), SC-514032 were supplied by Santa Cruz Biotechnology (Dallas, TX, USA). Mouse monoclonal anti-GAPDH, #AM4300, came from Thermo Fisher Scientific (Waltham, MA, USA). F-Actin was fluorescently labeled with TRITC-phalloidin from Sigma-Aldrich (St. Louis, MO, USA). Nuclei were labeled with DAPI staining (Sigma).

Secondary antibodies used in this study included: Goat anti-mouse IgG conjugated to Cy5 (Jackson ImmunoResearch Laboratories, West Grove, PA, USA); goat anti-rabbit IgG H&L (HRP) #ab97080; and goat anti-mouse IgG H&L (HRP) #ab97040 (Abcam, Cambridge Science Park, Cambridge, UK).

### Plasmids, siRNA and transfections

IRES-GFP empty vector, AXL-WT-IRES-GFP, and AXL-KD-IRES GFP were a kind gift from Dr. Aaron S. Meyer (Koch Institute for Integrative Cancer Research at MIT, Cambridge, MA, USA).

DNA transfection: Cells were transfected using Lipofectamine2000 (Invitrogen, Carlsbad, CA, USA) in antibiotic-free media, according to the manufacturer’s instructions. Media were replaced with full media after 6h. After 48 h, cells were replated onto cross-linked gelatin-coated plates (see Gelatin coating section, below) for 3-6h, depending on the experiment, then either fixed, or lysed for protein.

RNA transfection: Transfection of the siRNA siGENOME SMARTpool™ (GE Healthcare Dharmacon, Lafayette, CO, USA) was performed using DharmaFect1 (Dharmacon) at a 50 μM concentration. In every transfection, siNON-TARGETING POOL #2 was used as a control, with siTOX as the transfection efficiency reporter. Cells were incubated for 48 h before replating for an experiment. RNA sequences are listed in Supplementary Material 2.

AXL knockdown by shRNA: TRC AXL shRNA was purchased from GE Healthcare Dharmacon. pCMV-VSV-G (helper plasmid) and pHRCMV-8.2ΔR (packaging plasmid) were both kindly provided by Prof. Yardena Samuels (Weizmann Institute of Science, Rehovot, Israel). Mission pLKO.1-puro (Sigma-Aldrich) was used as a negative control vector. HEK293 cells were transfected with the target plasmid, together with the helper and the packaging plasmid, using Lipofectamine 2000 (Invitrogen). Conditioned medium was collected from the cells 48 h post-transfection, and placed on the target cells (A375) for 24 h. Cells were treated with 3μg/ml puromycin for selection. AXL knock-down was tested by real-time PCR, and was found to be 85% reduced (data not shown).

### Cell cultures

A375 metastatic melanoma cells, and HEK293, 4T1 and MDA-231 metastatic breast carcinoma cells were obtained from the American Type Culture Collection (ATCC) (Manassas, VA, USA). WM793 and A2508 cells were a kind gift from Prof. Meenhard Herlyn (Wistar Institute, Philadelphia, PA, USA). A subset of cell lines (104T, 63T and 76T) used in the study were derived from a panel of pathology-confirmed metastatic melanoma tumor resections collected from patients enrolled in Institutional Review Board (IRB)-approved clinical trials at the Surgery Branch of the National Cancer Institute (Bethesda, MD, USA). Pathology-confirmed melanoma cell lines were derived from mechanically or enzymatically dispersed tumor cells. Cells were cultured in DMEM or RPMI medium supplemented with 10% FCS (Gibco, Grand Island, NY, USA), 2 mM glutamine, and 100 U/mL penicillin-streptomycin. Cultures were maintained in a humidified atmosphere of 5% CO_2_ in air, at 37°C.

### Quantitative real-time PCR (QRT–PCR)

Total RNA was isolated using an RNeasy Mini Kit (Qiagen, Valencia, CA, USA), according to the manufacturer’s protocol. A 2 μg aliquot of total RNA was reverse-transcribed, using a high-capacity cDNA reverse transcription kit (Applied Biosystems, Carlsbad, CA, USA). Quantitative real-time PCR (QRT–PCR) was performed with a OneStep instrument (Applied Biosystems), using Fast SYBR® Green Master Mix (Applied Biosystems). Gene values were normalized to a GAPDH housekeeping gene. Primer sequences can be found in Supplementary Material 2.

### Gelatin coating

Gelatin gel: 96-well glass-bottomed plates (Nunc, Thermo Fisher Scientific) or plastic tissue culture plates were coated with 50 μg/ml poly-L-lysine solution (Sigma, catalogue # P-7405) in Dulbecco’s PBS, and incubated for 20 min at room temperature. The plates were then gently washed 3 times with PBS. Porcine skin gelatin, 0.2 mg (Sigma, catalogue #G2500) was dissolved at 37°C in 100 ml ddH_2_O. Gelatin was cross-linked with 1-ethyl-3-(3-dimethylaminopropyl) carbodiimide (EDC, Sigma, catalogue #03450) and N-hydroxysuccinimide (NHS, Sigma, catalogue #130672), each prepared as 10% solutions in ddH2O. The 96-well glass-bottomed plates were coated with 50 μl of gelatin and cross-linker mixture, and the plastic plates were coated with various volumes, according to their size. The ratios of gelatin to cross-linkers were 82.5:12.5:5 (gelatin: NHS: EDC).

Surfaces were incubated for 1 h at room temperature, washed 3 times in PBS, and sterilized by 30 min of UV radiation.

### Immunofluorescence staining, immunofluorescence microscopy, and image analysis

For immunostaining, cells were plated at ∼70% confluence on gelatin-coated glass-bottomed plates (see above) for varying time periods. Cells were then fixed for 3 min in warm 3% PFA (Merck, Darmstadt, Germany), 0.5% Triton X-100 (Fluka-Chemie AG, Switzerland), followed by 3% PFA alone for an additional 30 min. Post-fixation, cells were washed 3 times with PBS and incubated with primary antibody for 1 h, washed 3 times in PBS, and incubated for an additional 30 min with the secondary antibody, washed again, and kept in PBS for imaging.

Images were acquired using the DeltaVision microtiter system (Applied Precision, Inc., Issaquah, WA, USA), using a 40x/0.75 air objective, or a 60x/1.42 oil objective (Olympus, Tokyo, Japan). Image analysis was performed using ImageJ software (rsbweb.nih.gov/ij).

### Gelatin degradation assay

Fluorescently labeled gelatin (0.2 mg porcine skin gelatin; Sigma, catalogue #G2500) was prepared by Alexa Fluor™ 488 (A10235)/ Alexa Fluor™ 546 Protein Labeling Kit), molecular probes, Thermo Fisher Scientific, according to the manufacturer’s instructions.

Glass-bottomed 96-well plates were coated as mentioned above (gelatin coating), with ratios of 1:10 labeled gelatin: non-labeled gelatin.

For degradation assays, cells were plated on the gelatin matrix, and cultured for varying lengths of time (on average, 5-6h). Cells were fixed and stained for actin and DAPI, and degradation area was assessed by ImageJ software.

In every well, 36-64 fields of view were imaged in an automated fashion by the DeltaVision microtiter system, using a 40x objective. Cells in every field of view were counted using DAPI staining, and the degradation area was calculated by the fluorescently labeled gelatin channel, using the Analyze Particles plug-in in ImageJ software. The value for each well assessing invadopodia function was total degradation area/cell (μm^2^). To compare results between experiments, control values were normalized to 1, and all the samples were normalized accordingly.

For R428 inhibitor experiments, cells were plated on gelatin-coated plates with the indicated concentrations of the inhibitor (1-5 μM, depending on the cell type) or DMSO, and fixed after 5h. For prolonged R428 treatment, cells were cultured with the indicated concentrations of the inhibitor (1-2 μM, depending on the cell type) or DMSO for 2 days, then replated on gelatin-coated plates with the same inhibitor concentrations, and fixed (after 5h) for the invadopodia assay.

Invadopodia assay: Upon Gas6 stimulation, A375 cells were cultured with 200-1000 ng/ml of Gas6 for 5h. Cells were fixed and stained for actin and DAPI, invadopodia function was assessed as mentioned above, and the results compared to control cells. For 63T cells, Gas6 stimulation in concentrations of 100-1000 ng/ml was conducted under 1% serum conditions (only throughout the duration of the experiment).

### Immunoblotting

Whole-cell lysates were prepared using SDS lysis buffer (2.5% SDS, 25% glycerol, 125mM Tris-HCL pH=6.8, 0.01% bromophenol blue, and 4% freshly added beta-mercaptoethanol). Samples were resolved on 8% SDS-PAGE gels, and transferred into PVDF/nitrocellulose membranes. Blots were probed with antibodies to AXL (1:1000), p-ERBB3 (1:1000), ERBB3 (1:500), pCortactin (1:1000), cortactin (1:1000), pSRC (1:1000), Src (1:1000), pAKT (1:1000), AKT (1:1000), and GAPDH (1:1000) (see Antibodies and reagents section, above), and developed using SuperSignal® ECL reagents (Thermo Fisher Scientific).

For R428 inhibitor experiments, cells were plated on gelatin-coated plates with the indicated concentrations of the inhibitor (1-2 μM, depending on the cell type) or DMSO, and lysed for immunoblotting after 3h. For prolonged R428 treatment, cells were cultured with the indicated concentrations of the inhibitor (1-2 μM, depending on the cell type) or DMSO for 2 days, then replated on gelatin-coated plates with the same inhibitor concentrations, and lysed for immunoblotting after 3h.

### Cell invasion assay

A375 cells (50,000 cells) were cultured in serum-free media in the upper chamber of a BioCoat Matrigel Invasion Chamber #354480 (Corning, NY, USA), in duplicates Full medium containing 10% FBS was placed in the lower chamber. Cells were allowed to invade into the Matrigel for 24 h, then fixed with methanol, and stained with Gimsa. Cells that remained in the upper part of the chamber were removed with a cotton bud, and the inserts were washed 3 times, before imaging the inserts. Four quarters were imaged for each filter using a 10x lens, and cells were counted manually, using an ImageJ cell counter. Cell counts from the two duplicates were averaged. Each experiment was normalized to its control, as a percentage of invading cells.

### Screen targeting tyrosine kinome using siRNA

A screen for invadopodia regulators among tyrosine kinase family members was carried out, using a tyrosine kinase library (siGENOME SMARTpool™, Dharmacon) for A375 and 63T metastatic melanoma cells (see Cell cultures section, for details of cell lines).

The siRNA library targets 85 genes of the tyrosine kinome (See Supplementary Material 1). Cells were cultured in a 96-well plate format in antibiotic-free media, 1 day prior to transfection. The next day, the siRNA library was transfected into the cells, using a DharmaFect1 protocol calibrated to achieve the highest transfection efficiency in those cells. Cells were transfected with 50µM siRNA that targets tyrosine kinases, using non-targeting siRNA as a control (Supplementary Fig. 1A). In each experiment, siTOX was used to test transfection efficiency in a qualitative fashion (Fig. 1B, 1F).

In order to test transfection efficiency in a quantitative fashion, a few wells in every experiment were transfected with siCON (non-targeting control) or with siSRC, and Src levels were analyzed using real-time PCR (Fig. 1C, 1E). Transfection efficiency was 80-90%. The following day, the medium was replaced by full media, and cells were incubated for another 48 h. Three days following transfection, cells were replated in fluorescently labeled gelatin on 96-well glass-bottomed plates (see Gelatin coating section, above). After 6 h, cells were fixed and stained for actin, TKS5 and DAPI. Plates were then imaged in an automated fashion with a DeltaVision microtiter system, using a 40x objective. In each well, 16-36 fields of the gelatin, actin, TKS5 and DAPI were imaged automatically by the microscope; each field was then analyzed using ImageJ software for degradation/cell (µm^2^), and all the fields were quantified into average degradation rates per cell (Supplementary Fig. 1B).

Quantities and shapes of invadopodia were analyzed in a qualitative and quantitative fashion, using actin and TKS5 staining (Table 1, Phenotype; Supplementary Material 1; and Supplementary Fig. 1B, 1C).

In Supplementary Material 1, data from the 3 screens in the A375 cell line, and from the 2 screens in the 63T cell line, are presented. In every tab, control values are presented first, followed by all the genes that were analyzed. Degradation values were normalized to control, that was set as 1, in order to compare different experiments. A *p*-value was calculated for each gene, according to the degradation/cell values in all the fields, relative to the control values. In the first screen, only knock-down of genes that showed any effect on matrix degradation (52 of 85 tyrosine kinases) were analyzed. Genes whose knock-down caused a statistically significant difference (*p* value ≤ 0.05) and led to a ≤ 3-fold reduction or a ≥ 1.5-fold elevation or more in gelatin degradation, were chosen for further validation (Supplementary Material 1, Screen 2). Altogether, the primary screen was performed three times for the final hits (See results in Supplementary Material 1).

In 63T cells, the 10 hits which were found in A375 cells were tested two additional times (Supplementary Material 1).

### Invadopodia quantification

Invadopodia quantification was percentage of invadopodia forming cells in 16-36 fields, based on their actin and TKS5 labeling. Cell count was performed by DAPI, as described above.

### Gene expression analysis

Gene expression profiles of AXL, ERBB3, MITF, and three of the MITF targets (PMEL, TYRP1, and MELANA), were extracted from 62 melanoma cell lines from the *CCLE* dataset (https://portals.broadinstitute.org/ccle) containing Affymetrix microarray expression data. Hierarchical clustering by Euclidian distance was applied on the Z-score values, and four groups of cell lines were manually defined. *P*-values were calculated for the differences between ERBB3 and AXL levels, and the other 4 genes within each group (see Supplementary Material 3, tab 1). Invadopodia-related genes and tyrosine kinases were compared only between two groups of cell lines with alternating patterns of AXL and ERBB3 (Clusters 1 and 2) (see Supplementary Material 3, tabs 2 and 3). Significance level was determined by a Mann-Whitney two-tailed test, following multiple hypothesis correction.

### Statistical analysis

Quantitative data for statistical analysis were expressed as mean ± SEM (shown as an error bar) from at least three independent experiments, unless stated otherwise. The significance of the difference between samples was calculated using a two-tailed Student’s t-test. *P*-value > 0.05 - n.s (not significant); *p*-value ≤ 0.05 *; *p*-value ≤ 0.01 **; *p*-value ≤ 0.001 ***; *p*- value ≤ 0.0001 ****.

## Supplementary Figures

**Figure S1:**
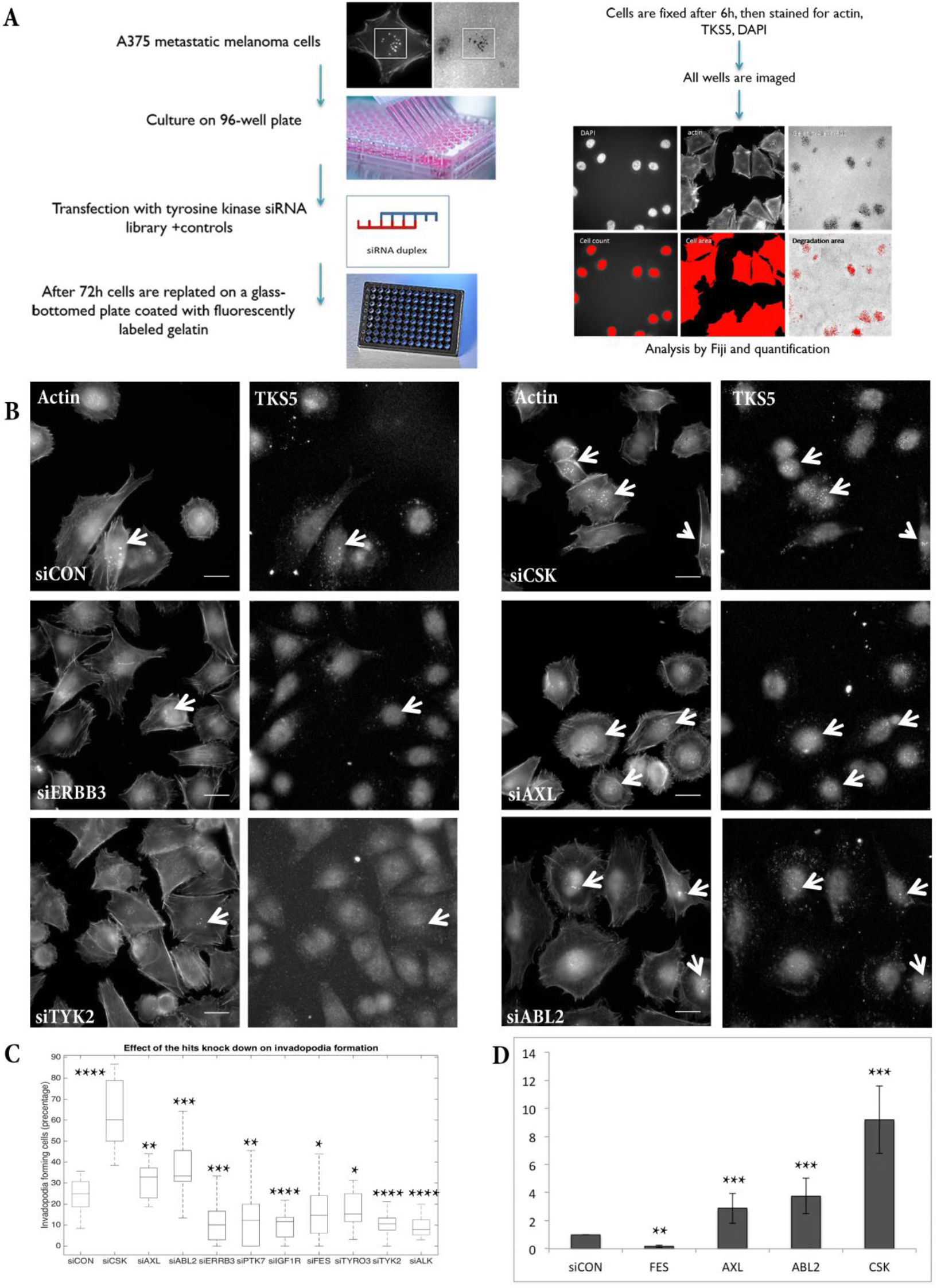
Screen workflow and results. (A) Cells were cultured on 96-well plates, 1 day prior to transfection. Cells were transfected with a tyrosine kinase siRNA library (Dharmacon), using the DharmaFECT 1 transfection reagent. Following 72 h transfection, cells were replated on fluorescently labeled, gelatin-coated 96-well plates for 6 h. Cells were fixed and stained for actin, TKS5 and DAPI, then imaged and analyzed using ImageJ software. Quantification was measured as degradation area/cell (μm^2^). (B) Representative images from the A375 cell screen labeled with actin and TKS5, to demonstrate invadopodia enrichment or reduction upon knockdown of specific genes. White arrows denote invadopodia. Scale bar: 20 μm. (C) Quantification of invadopodia-forming cells (expressed as a percentage) upon knock-down of the hit genes in A375 cells. (D) Quantification of normalized degradation/cell for the final hits of the screen (invadopodia assay) in 63T cells. Two repeats; n>100 in each sample.

**Figure S2:**
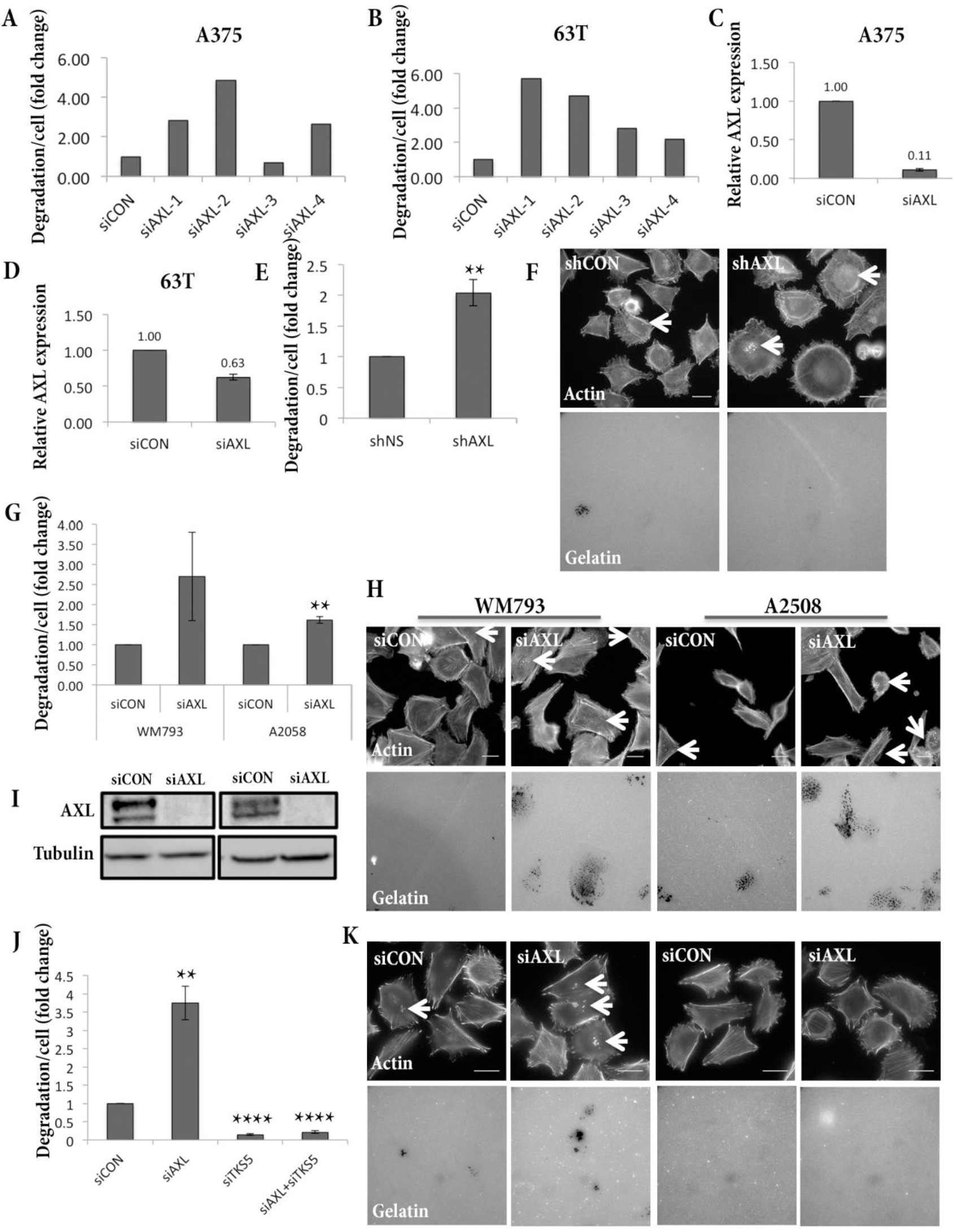
AXL regulates invadopodia structures in melanoma cells. (A, B) Invadopodia assay of A375 (A) or 63T cells (B) transfected with siCON or 4 individual siRNA-targeting AXL, shows that the phenotype of invadopodia degradation elevation repeats in 3 out of 4 siRNA duplexes. Two repeats for each cell; n>100 in each sample. (C, D) Real-time analysis of AXL gene expression in siCON *versus* siAXL A375 cells (C) or 63T cells (D), shows the reduction in AXL expression levels upon knock-down; n=3 for each cell. (E, F) Invadopodia assay of A375 cells stably expressing shRNA targeting AXL or shCONTROL, cultured on fluorescently labeled gelatin for 6 h, shows that invadopodia elevation upon AXL knockdown also occurred using AXL shRNA. Three repeats; n>100 in each sample. Scale bar: 20 μm. Cells were stained for actin. (G, H) Invadopodia assay of siCON *versus* siAXL in WM793 and A2508 melanoma cells. Cells were cultured on fluorescently labeled gelatin for 6 h. Two repeats; n>100 in each sample. Scale bar: 20 μm. (I) Western blot analysis of AXL expression in siCON versus siAXL WM793 and A2058 cells. Tubulin was used as a control. (J, K) Invadopodia assay of A375 cells with knock-down of AXL, TKS5, or AXL+TKS5 versus control, cultured on fluorescently labeled gelatin for 6 h. Cells were stained for actin. Three repeats; n>100 in each sample. Scale bar: 20 μm.

**Figure S3:**
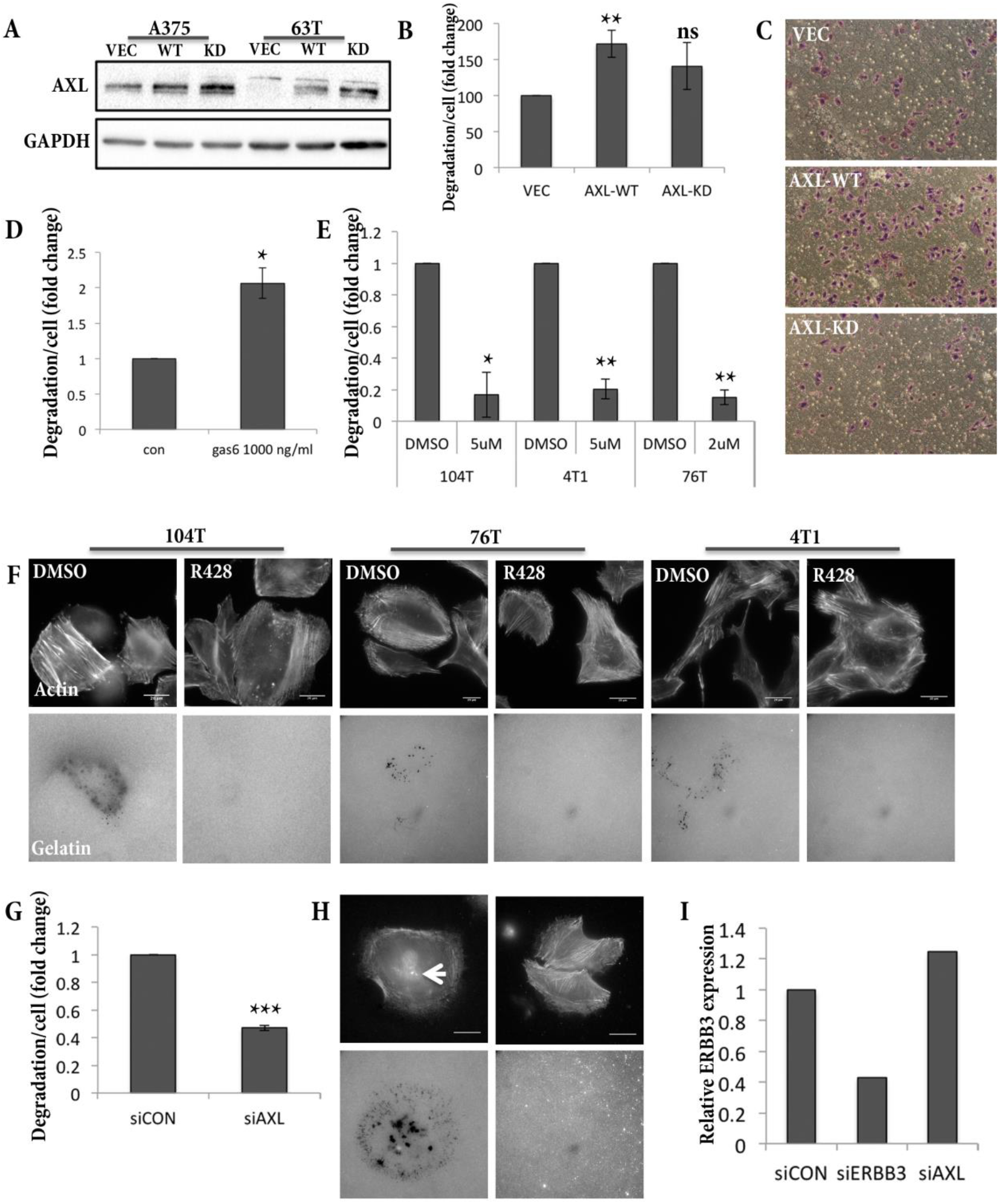
AXL is a positive regulator of invadopodia. (A) AXL Western blot analysis of A375 and 63T cells overexpressing AXL-WT, AXL-kinase dead (AXL-KD), or control vector, shows elevated AXL levels upon overexpression. GAPDH was used as a loading control. (B) Trans-well invasion assay of A375 cells overexpressing AXL-WT, AXL-kinase dead (AXL-KD), or control vector; n=3. (C) Representative images of the invasion inserts in (B). (D) Invadopodia assay of control or 1000ng/ml Gas6 63T-treated cells; cells were cultured on fluorescently labeled gelatin for 6 h. Stimulation was performed in 1% serum. Three repeats for each cell; n>100 in each sample. Scale bar: 20 μm. (E, F) Invadopodia assay of 104T, 4T1 and 76T cells cultured on fluorescently labeled gelatin for 6 h with DMSO, or the AXL inhibitor R428 (in concentrations of 5μM, 5μM and 2μM, respectively). Cells were stained for actin. Three repeats for each cell; n>100 in each sample. Scale bar: 20 μm. (G, H) Invadopodia assay of siCON versus siAXL MDA-231 cells cultured on fluorescently labeled gelatin for 6 h, shows a reduction in invadopodia function upon AXL knock-down. Three repeats; n>100 in each sample. Scale bar: 20 μm. (I) Real-time analysis of ERBB3 gene expression in siCON, siAXL or siERBB3 A375 cells shows a reduction in ERBB3 expression upon KD, and a mild elevation in ERBB3 expression upon AXL knockdown; n=3.

**Figure S4:**
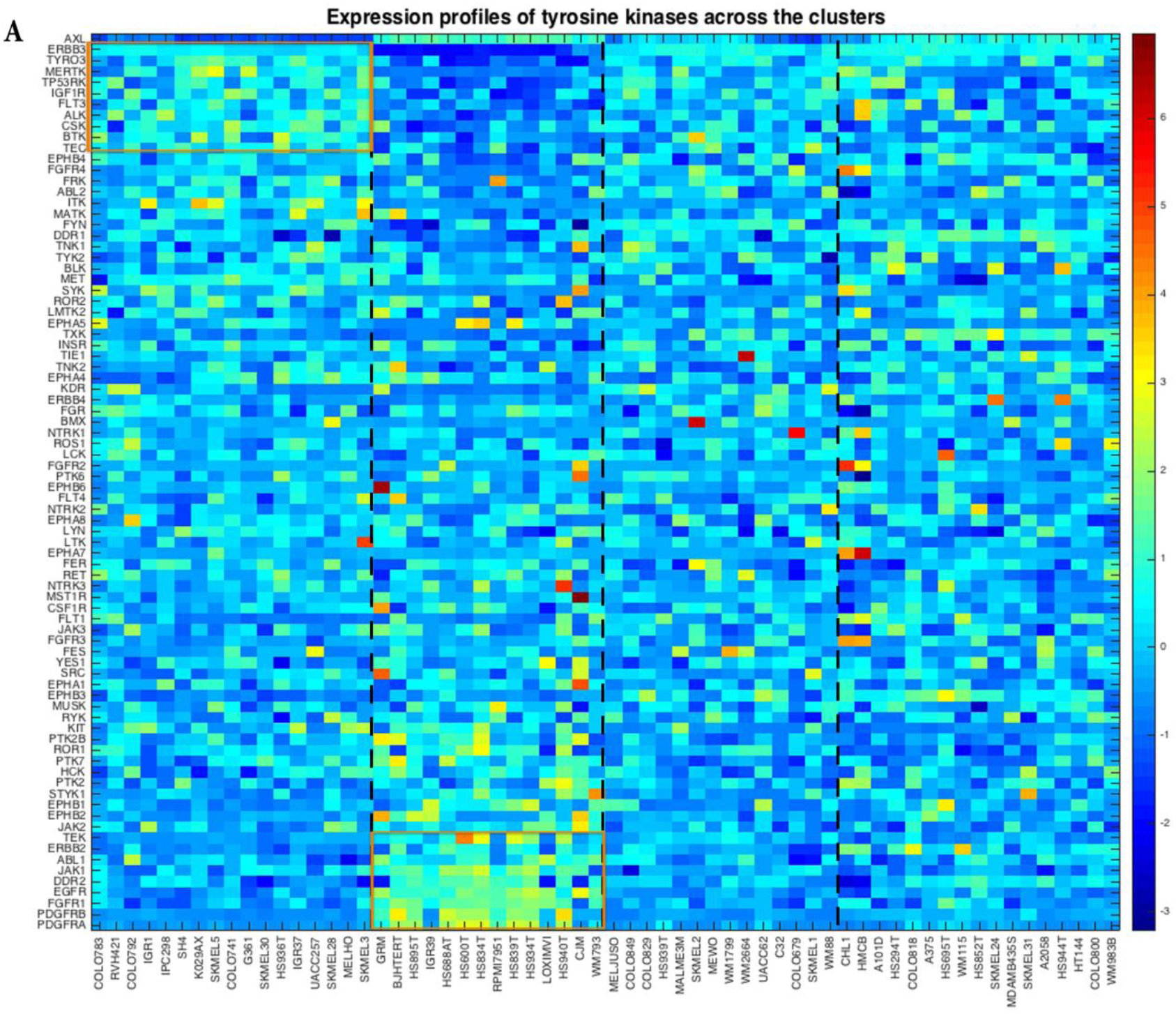
Analysis of tyrosine kinome and invadopodia components in the CCLE melanoma database. Analysis of Z-score values of 80 tyrosine kinases across the 4 CCLE clusters presented in Fig. 6A. Significance of differences in TK expression between Clusters 1 and 2 are presented in Supplementary Material 3, tab 3.

## Supplementary Material

**Supplementary Material 1:** In the Excel file, the raw data for the screens (3 in A375 cells, and 2 in 63T cells) is contained under every tab, the data for each screen are quantified, and the results, summarized.

**Supplementary Material 2**: A list of primers and siRNA sequences utilized in the study.

**Supplementary Material 3:** Statistics for gene expression analysis. In the first tab, significant levels of ERRB3 and AXL expression against all 6 genes (AXL, ERBB3, MITF3 and the 3 targets) in each cluster. In the second tab, significant levels of differences in invadopodia genes between Clusters 1 and 2, as well as the delta median of expression for each gene. In the third tab, significant levels of the differences in TK genes between Clusters 1 and 2, as well as the delta median of expression for each gene.

## List of Abbreviations

RTK: Receptor Tyrosine Kinase
SFK: Src Family Kinase
MMPs: Metalloproteinases
ECM: Extracellular Matrix
KD: Knock-down

